# *SuSiE 2.0*: improved methods and implementations for genetic fine-mapping and phenotype prediction

**DOI:** 10.1101/2025.11.25.690514

**Authors:** Alexander McCreight, Yanghyeon Cho, Ruixi Li, Peter Carbonetto, Daniel Nachun, Hao-Yu Gan, Deborah Kunkel, Matthew Stephens, William R.P. Denault, Gao Wang

## Abstract

*SuSiE 2.0* is a redesign of `susieR` for statistical fine-mapping in genetics. Its shared fitting algorithm handles individual-level data or summary statistics paired with LD matrices or genotype sketches. We used this algorithm to implement several *SuSiE* extensions, including *SuSiE-inf, mvSuSiE, mfSuSiE*, and *SuSiE-rRSS*, speeding up *SuSiE-inf* by more than 30-fold in some benchmarks. In simulations with polygenic backgrounds, *SuSiE-inf* substantially lowered the credible set false discovery rate relative to *SuSiE*, with some loss of power. Across three ROSMAP brain regions, *SuSiE* and *SuSiE-inf* shared 75% of signals detected by either method and were the two most accurate predictors of gene expression.

## Background

The Sum of Single Effects (*SuSiE*) model [1] has emerged as a powerful approach for Bayesian variable selection, producing posterior inclusion probabilities (PIPs) and single-effect credible sets (CSs) that quantify uncertainty in the selected variables. These make *SuSiE* particularly suited for genetic fine-mapping, where causal signals are sparse yet genetic variables are highly correlated due to linkage disequilibrium (LD), and identifying candidate causal variants is essential for downstream biological interpretation. The `susieR` package has received over 560,000 downloads across Anaconda and CRAN (as of November 2026), and has inspired a substantial body of methodological extensions [2–16], with integration into major analysis pipelines including *coloc* [17] and large-scale analyses of UK Biobank [18], GTEx [19], and FinnGen [20].

However, the original `susieR` implementation suffers from architectural limitations that hinder extensibility and performance. The codebase lacks modular separation between data handling and core algorithm logic, requiring extensive changes to core functions for even modest methodological extensions. Meanwhile, many extensions exist only as standalone command-line tools [2, 7, 16, 21], making them difficult to integrate into R-based production pipelines or benchmark systematically.

Here we present *SuSiE 2.0*, a modular redesign of `susieR` that separates data handling from the shared fitting algorithm, which makes it easier to extend *SuSiE*. We reimplemented *SuSiE-inf* in `susieR` to support individual-level data alongside summary statistics, the only input type supported by the original Python implementation [2]. We also added a residual variance prior for small-sample calibration with both individual-level data and summary statistics [8]. Using the same design, we reimplemented *mvSuSiE* for joint fine-mapping of multiple traits [9] and *mfSuSiE* for fine-mapping of spatially correlated functional outcomes [10]. We also implemented *SuSiE-rRSS* for robust fine-mapping from summary statistics when reference LD is uncertain [22]. The redesign speeds up *SuSiE-inf* and fine-mapping from summary statistics (*SuSiE-RSS*), making both more practical for genome-wide analyses.

Polygenic backgrounds can inflate the false discovery rate (FDR) in fine-mapping [2]. Modeling these backgrounds can reduce false discoveries at the expense of power. In regions with multiple causal effects, the signals identified by a fit can also depend on its initialization and the LD among variants. We compare the signals reported by *SuSiE* and *SuSiE-inf* in simulations and in expression quantitative trait locus (eQTL) data. We also evaluate *SuSiE-ash*, which uses adaptive shrinkage to model background genetic effects, as a follow-up method for selected loci (**Methods**). We benchmark the gene expression predictions of *SuSiE* and *SuSiE-inf* against established methods for transcriptome-wide association studies (TWAS).

## Results and discussion

*SuSiE 2.0* uses S3 dispatch in the R programming language to make it straightforward to add new models and improve existing methods within a shared fitting algorithm (**Figure 1A**) [23]. The algorithm works with individual-level data and summary statistics, and extensions can change how the single effect regression (SER) computes Bayes factors or how the model objective is evaluated [1]. For example, *SuSiE-rRSS* adjusts the residual variance in the SER to account for finite reference sampling and differences in LD between the study and reference populations [22]. For fine-mapping from summary statistics, the same interface accepts either an LD matrix or a genotype sketch, a compact representation of LD [24]. We use standard *SuSiE* to test extensions in settings where the models coincide. With one scalar outcome and matched model settings, our tests check that *mvSuSiE* and *mfSuSiE* agree numerically with standard *SuSiE*.

**Fig. 1:**
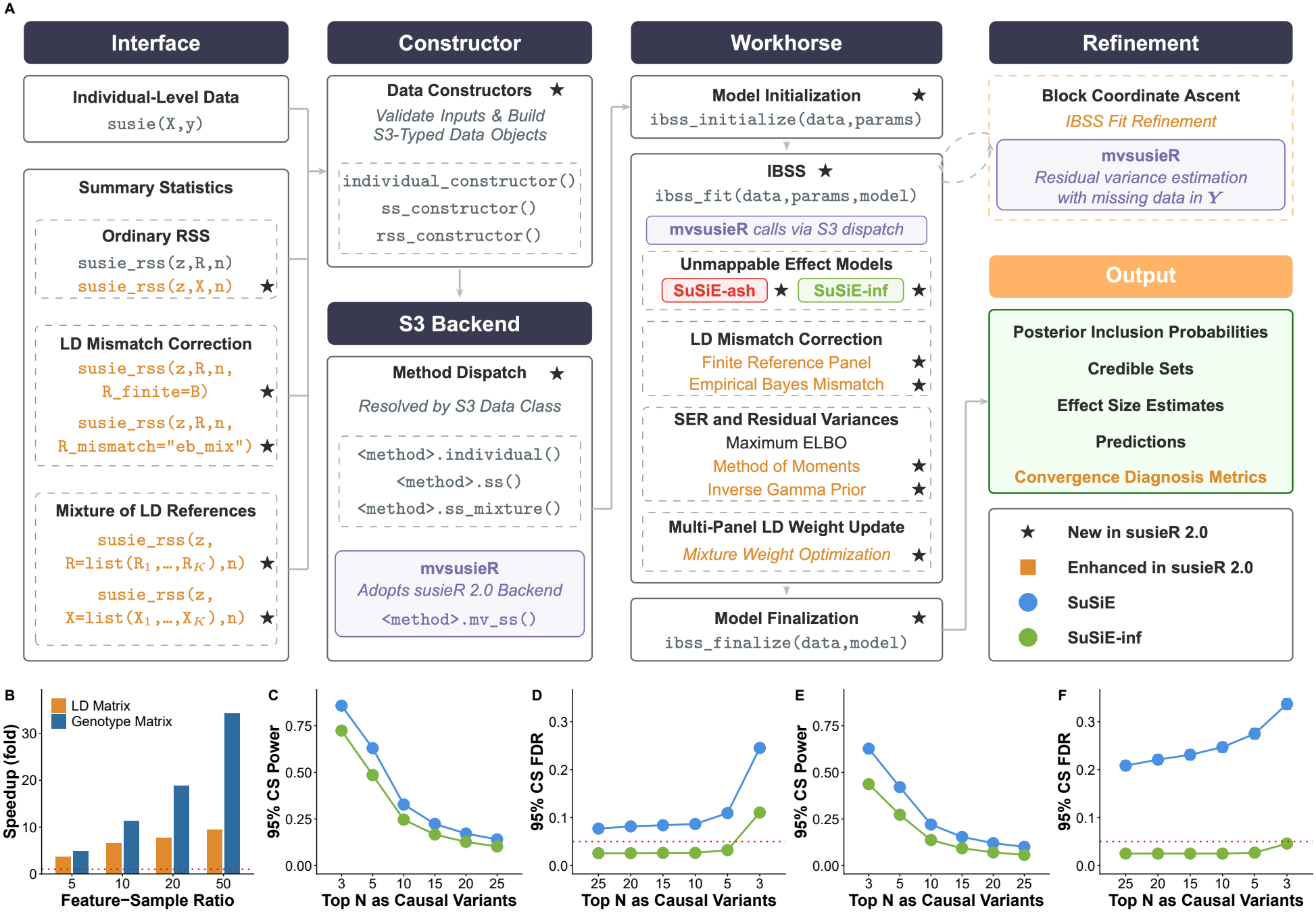
SuSiE 2.0 architecture and performance. (A) Modular architecture of *SuSiE 2.0* with S3 generic dispatch. (B) Fold speedup of *SuSiE-inf* over its original eigendecomposition-based implementation, for the summary-statistics (LD-matrix, orange) and individual-level (genotype-matrix, blue) backends, across variant-to-sample ratios (*p/n*) at *p* = 5,000. (C, E) 95% CS power and (D, F) 95% CS FDR for *SuSiE* and *SuSiE-inf*, as the number of top variants treated as causal varies, under oligogenic effects on (C, D) a moderate infinitesimal background and (E, F) an extensive infinitesimal background.

To accelerate *SuSiE-inf*, we exploit the low-rank structure of genotype-derived LD matrices when variants outnumber samples. For individual-level data, we compute a thin singular value decomposition (SVD) directly from the genotype matrix. For summary statistics, we use pivoted Cholesky factorization followed by a thin SVD of the resulting factor. These computations avoid the full LD eigendecomposition and, in our benchmarks, give results that agree numerically with those of the full eigendecomposition. **Figure 1B** summarizes the fold speedup of *SuSiE-inf* over its original implementation, with further gains when genotype matrices are used instead of LD matrices (**Methods** and **Supplementary Notes S5**). We separately optimized the summary statistics backend for the regularized RSS model [25], replacing dense matrix constructions that were rebuilt across iterations with algebraically equivalent matrix-vector operations. This gives a roughly 8-fold speedup on identical inputs, rising to 17-fold at the largest variant-to-sample ratio we tested when a genotype matrix replaces the dense LD matrix (**Supplementary Notes S6** and **Supplementary Figure S1**). *SuSiE 2.0* also provides detailed diagnostics, convergence monitoring, a block coordinate ascent refinement procedure that explores alternative initializations to improve the fit, and residual variance estimation with missing data in multi-trait fine-mapping. *SuSiE 2.0* maintains full backward compatibility with a comprehensive suite of unit tests.

We evaluated *SuSiE* and *SuSiE-inf* across diverse genetic architectures using *simxQTL*, an R package we developed for benchmarking molecular QTL studies that simulates realistic eQTL architectures [26] (**Methods**). In sparse simulations using typical 1-Mb *cis*-windows with realistic LD, *SuSiE* had the highest power among the methods compared and showed slightly inflated FDR. *SuSiE-inf* controlled FDR well with some loss of power (**Supplementary Figure S2**). In the more complex simulations, *SuSiE-inf* had substantially lower FDR than *SuSiE*, with lower power (**Figure 1C-F**). Under the extensive infinitesimal background, credible sets averaged 13.1 variants for *SuSiE-inf* and 31.8 for *SuSiE*, with median purity of 0.98 and 0.94, respectively (**Supplementary Figure S3**). For TWAS, *SuSiE* had a slight advantage in sparse simulations, and *SuSiE-inf* performed better as the number of signals increased (**Supplementary Figure S4**).

Under polygenic architectures, non-causal variants can produce spurious signals through LD with multiple causal variants (**Supplementary Figure S5**). *SuSiE-inf* can reduce these false discoveries by modeling the polygenic background with random effects, which can also absorb some sparse signals. In the complex simulations, *SuSiE-ash* had similar power to *SuSiE* and an FDR between those of *SuSiE* and *SuSiE-inf* (**Supplementary Figure S6**). At the simulated locus in **Supplementary Figure S5**, *SuSiE* reported four false-positive CSs. *SuSiE-inf* reported one true signal and *SuSiE-ash* reported three, with neither method reporting a false-positive CS.

To report credible sets without LD-based purity filtering, we explored two recently proposed methods for post-processing *SuSiE* fits, attainable coverage [21] and Bayesian Linear Programming (BLiP) [27] (**Supplementary Notes S9**). In our simulations, attainable coverage sometimes produced diffuse credible sets to reach the required 95% posterior mass. We therefore recommend using it only when LD is unavailable for purity filtering. BLiP offered limited gains in the scenarios we evaluated, so we did not include it in the current release (**Supplementary Figure S7**). *SuSiE 2.0* additionally offers a correlation-based credible set extension, which augments each CS with variants nearly indistinguishable from its members and gave a small but consistent improvement in both power and FDR across every architecture we tested (**Supplementary Notes S7** and **Supplementary Figures S8** and **S9**).

We applied *SuSiE* and *SuSiE-inf* to eQTL fine-mapping in three brain tissues from ROSMAP samples across 2,654 genes (**Methods**). Both methods yielded a similar median CS size (7.0 and 6.5) and purity (≥ 0.97). *SuSiE* identified more CSs than *SuSiE-inf* (4,059 versus 3,474), consistent with our simulations. CSs containing more than 10 variants accounted for just over half of this difference (308 of 585 CSs; **Figure 2A**).

**Fig. 2:**
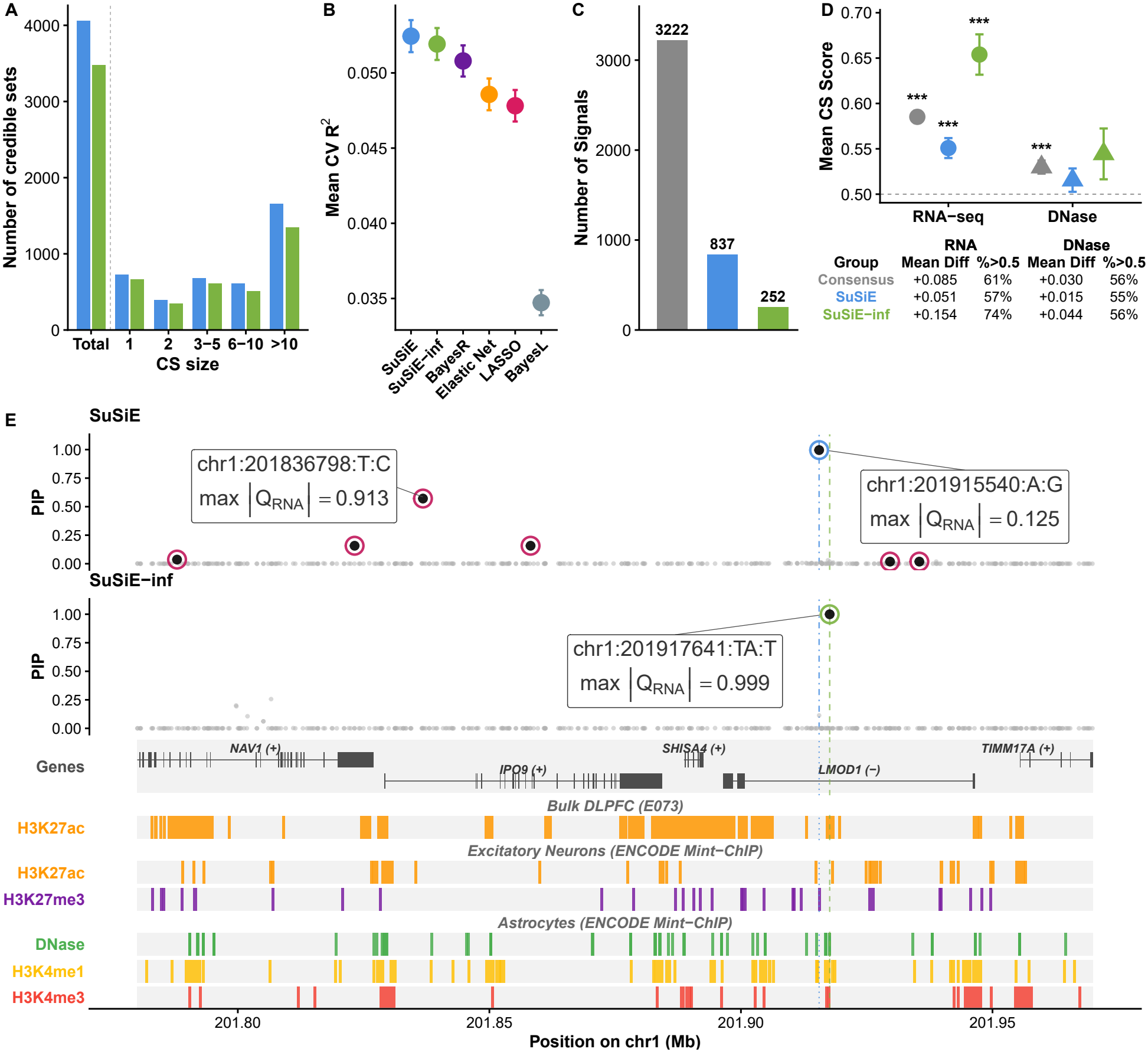
Fine-mapping and gene expression prediction in ROSMAP. (A) Number of credible sets identified by *SuSiE* and *SuSiE-inf*, in total and stratified by credible set size. (B) Mean cross-validated *R*^2^ for gene expression prediction methods. (C) Number of signals recovered by both methods, or unique to each. (D) PIP-weighted AlphaGenome functional enrichment scores by cross-method group, for RNA-seq (circles) and DNase (triangles), restricted to complex regions where *SuSiE* reports two or more CSs; points show group means, error bars show *±* 1 standard error, and asterisks mark means significantly above the 0.5 reference value in one-sided *t*-tests after Benjamini-Hochberg correction (adjusted *P* : ^*^ *<* 0.05, ^**^ *<* 0.01, ^***^ *<* 0.001); the table gives the mean difference from the 0.5 reference value and the percentage of signals exceeding it. (E) Fine-mapping at the *LMOD1* locus, showing per-method PIPs and tissue-matched regulatory annotation tracks.

To assess predicted functional support for the fine-mapped signals, we obtained tissuematched regulatory predictions from AlphaGenome [28] and computed a PIP-weighted functional enrichment score for each CS. We matched CSs across methods by Jaccard overlap (**Methods**). Both methods recovered 3,222 of 4,311 distinct signals, with 837 unique to *SuSiE* and 252 to *SuSiE-inf* (**Figure 2C**). Among loci where *SuSiE* reported at least two CSs, the mean RNA-seq score in each of the three groups was significantly above the reference value of 0.5 after FDR correction. *SuSiE-inf*-unique signals had the highest mean score (0.654, with 74% above 0.5), followed by consensus signals (0.585, 61%) and *SuSiE*-unique signals (0.551, 57%). DNase-based scores reached significance only for the consensus group (**Figure 2D**).

At the *LMOD1* locus, *SuSiE* and *SuSiE-inf* report three CSs in total, with no matches across methods (**Figure 2E**). *SuSiE* prioritizes chr1:201915540:A:G (PIP 0.996), and *SuSiE-inf* prioritizes chr1:201917641:TA:T (PIP 0.9996). The variants are 2.1 kb apart and have *r*^2^ = 0.37. The maximum absolute RNA scores in the corresponding credible sets were 0.999 for *SuSiE-inf* and 0.125 for *SuSiE*. The variant prioritized by *SuSiE-inf* overlaps bulk H3K27ac and astrocyte DNase, H3K4me1 and H3K4me3 peaks consistent with activating regulatory elements. The variant prioritized by *SuSiE* overlaps excitatory neuron H3K27me3, a repressive histone mark. *SuSiE* also reports a diffuse CS of six variants spanning roughly 150 kb. Its lead variant is chr1:201836798:T:C (PIP 0.571), and the maximum absolute RNA score across the CS was 0.913. In a follow-up comparison at this locus, *SuSiE-ash* reported an additional credible set near chr1:201806618 (**Supplementary Figure S10**).

In ROSMAP, we compared *SuSiE* and *SuSiE-inf* against other commonly used gene expression prediction methods [29–33] and found that *SuSiE* achieved the highest mean cross-validated *R*^2^ (**Figure 2B**). These results support using *SuSiE*-based methods to construct expression predictors for TWAS.

## Conclusions

For genome-wide scans, we recommend *SuSiE* when speed and power are the priority, accelerated *SuSiE-inf* when false discovery control is a priority or substantial polygenic background effects are expected, or both to compare the signals they report (**Figure 1B-F** and **Supplementary Figure S11**). For follow-up at selected loci, *SuSiE-ash* can optionally be used to assess sensitivity to modeling and fitting choices and investigate additional candidate signals. The same fitting code that supports functional regression for chromatin accessibility profiles across cell types in *mfSuSiE* [10] would make it easier for developers to add *SuSiE* linear regression and analyze these profiles jointly with gene expression and splicing. *SuSiE 2.0* and the companion simulation package *simxQTL* are available as documented R packages.

## Methods

### Accelerating *SuSiE-inf*

The original *SuSiE-inf* implementation [2] performs a full eigendecomposition of the LD matrix at a one-time cost of *O*(*p*^3^) in the number of variants *p* regardless of sample size. Because the LD matrix is low-rank when variants outnumber samples, as in most QTL settings, we replace the full eigendecomposition with cheaper low-rank factorizations. When only an LD matrix is available, we use a pivoted Cholesky factorization followed by a singular value decomposition (SVD), and when individual-level data are supplied, we take a thin SVD of the genotype matrix directly. In our benchmarks, both approaches gave results that agreed numerically with those of the original implementation. Implementation details, complexity, and numerical validation are provided in **Supplementary Notes S5**.

### Overview of *SuSiE-ash* Model

*SuSiE-ash* extends *SuSiE* by jointly modeling sparse causal effects ***β*** via the Sum of Single Effects framework and unmappable background effects ***θ*** via an adaptive-shrinkage mixture-of-normals prior implemented through *Mr.ASH* [29]. Posterior inference proceeds by coordinate-ascent variational inference, alternating between single-effect regression updates for ***β*** and normal-means updates for ***θ***. A diffusion-aware effect separation mechanism coordinates these updates based on credible set purity and cross-effect collision history, selectively subtracting well-localized sparse effects from *Mr.ASH* residuals while protecting effects still under resolution from being absorbed into the background component. Model specification, variational updates and diffusion-aware effect separation are described in **Supplementary Notes S1-S3**.

### Overview of Simulation Study Design

We evaluated *SuSiE, SuSiE-ash*, and *SuSiE-inf* using UK Biobank genotype data under two settings: sparse architectures with 1-5 causal variants and fixed per-SNP heritability, and complex architectures partitioning genetic effects into sparse, oligogenic, and polygenic (or infinitesimal) components with total *h*^2^ = 0.25. For complex architectures, we evaluated CS power and FDR relative to the variants with the largest absolute nonzero simulated effects, varying the number *N* included in this target set. CSs containing none of these variants were counted as false discoveries. Simulation designs are implemented in *simxQTL*, an R package for benchmarking of gene-mapping methods (GWAS and xQTL, https://github.com/StatFunGen/simxQTL). See **Supplementary Notes S8** for full details.

### Overview of eQTL Data

We used bulk expression quantitative trait loci (eQTL) data from three brain cortical regions: dorsolateral prefrontal cortex (DLPFC; *n* = 784), anterior cingulate cortex (AC; *n* = 593), and posterior cingulate cortex (PCC; *n* = 441), derived from post-mortem tissue of Religious Orders Study and Memory and Aging Project (ROSMAP) donors [34]. We followed the same preprocessing steps as described in Cao et al. [35].

### Functional Enrichment of Credible Sets

We scored variants within 95% credible sets using tissue-matched AlphaGenome predictions and defined a PIP-weighted CS score to quantify functional relevance. To compare methods in complex regions, we restricted analysis to loci where *SuSiE* identified two or more CSs and classified CSs from *SuSiE* and *SuSiE-inf* into three groups based on cross-method concordance, defining matching CSs by a Jaccard index ≥ 0.75. For each matched signal, we averaged the constituent CS scores separately for RNA-seq and DNase. Significance was assessed via one-sided *t*-tests with Benjamini-Hochberg FDR correction. Full details are in **Supplementary Notes S10**.

## Declarations

### Ethics approval and consent to participate

The Religious Orders Study, Rush Memory and Aging Project, and Rush Alzheimer’s Disease Center Repository were approved by the Institutional Review Board of Rush University Medical Center (protocols L91020181, L86121802, and L99032481, respectively). ROSMAP participants provided written informed consent. Columbia University Institutional Review Board 5 approved molecular profiling and secondary analyses for the FunGen-xQTL project (protocol IRB-AAAR4962). The UK Biobank Resource received ethics approval from the North West-Haydock Research Ethics Committee, and participants provided informed consent. We used the UK Biobank Resource under application 27386.

### Consent for publication

Not applicable.

### Availability of data and materials

ROSMAP eQTL summary statistics are available through the FunGen-xQTL Atlas at NIA-GADS (NG00184.v1, https://dss.niagads.org/datasets/ng00184/) and can be queried at https://xqtl.niagads.org/. Access to individual-level ROSMAP whole-genome sequencing data is managed by NIAGADS (NG00067, https://dss.niagads.org/datasets/ng00067/). Access to the UK Biobank genotypes used in the simulations is managed by UK Biobank (https://www.ukbiobank.ac.uk/use-our-data/apply-for-access/).

### Code availability

*SuSiE 2.0* is available in the *susieR* package (https://stephenslab.github.io/susieR) under the BSD 3-Clause license. The source for version 2.0.0 is at https://github.com/stephenslab/susieR/tree/2.0.0. The package is written in R, with C++ routines for *SuSiE-ash*. It requires R 3.0.2 or later and a C++ toolchain for installation from source. New features are summarized at https://stephenslab.github.io/susieR/articles/announcements.html. The *mvsusieR* (https://stephenslab.github.io/mvsusieR/) and *mfsusieR* (https://statfungen.github.io/mfsusieR/) packages use the *SuSiE 2.0* infrastructure. Simulation functions are provided in the *simxQTL* package (https://statfungen.github.io/simxQTL). Real-data analysis pipelines (fine-mapping and TWAS) are available at https://github.com/StatFunGen/xqtl-protocol and https://github.com/StatFunGen/pecotmr. Analysis and figure scripts are available at github.com/alexmccreight/susieR2.0-paper.

### Authors’ contributions

AM developed the software architecture of *SuSiE 2.0* with input from GW and implemented the numerical experiments. YC and AM developed the details of the *SuSiE-ash* model. RL and AM performed the data analyses. PC, DN and HG contributed to improvements in the *SuSiE 2.0* package. DK contributed to the summary statistics backend of *SuSiE-ash*. AM, YC and GW wrote the manuscript. GW conceived the project and its methods and designed the experiments. MS, WD and GW jointly supervised the research.

## Acknowledgements

We thank the participants and investigators of the Religious Orders Study and the Rush Memory and Aging Project (ROSMAP).

## Supplementary Notes

### S1 *SuSiE-ash* Model and Motivation

*SuSiE-ash* is a Bayesian variable selection approach that improves fine-mapping by combining *SuSiE* [1] for sparse variable selection and *Mr.ASH* [29] for adaptive shrinkage estimation of unmappable effects. The key idea is to iteratively update the sparse effects ***β*** using *SuSiE* on residuals that account for ***θ***, followed by updating the unmappable effects ***θ*** using *Mr.ASH* on residuals after removing well-localized sparse effects.

*SuSiE-ash* is based upon the following model:

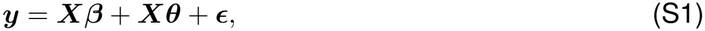

Here ***y*** is the centered phenotype vector for *n* individuals, and ***X*** = [***x***_1_, …, ***x***_*p*_] is the standardized *n* × *p* genotype matrix for *p* variants in the region. The sparse effects ***β*** and unmappable effects ***θ*** are independent *p*-vectors. The residual is ***ϵ*** ~ *N* (**0**, *σ*^2^***I***).

Following standard *SuSiE* [1], we write the sparse component ***β*** as a sum of *L* single effects. More than one single effect can select the same variant.

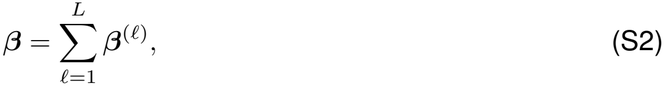

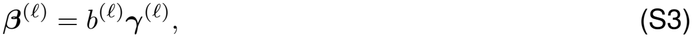

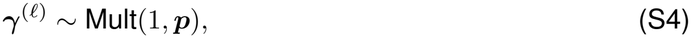

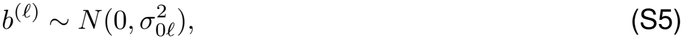

where ***β***^(*l*)^ denotes the *l*-th single-effect vector, 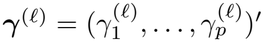 is a *p*-vector indicating the location of the causal SNP in the *l*-th single effect, with ***p*** = (*p*_1_, …, *p*_*p*_)^*′*^ representing the prior weights that sum to 1, and *b*^(*l*)^ is a scalar representing the causal effect size in the *l*-th single effect. When *L* = 1, the sparse component has the single-effect regression (SER) form.

We construct ***θ*** using an adaptive shrinkage prior for the scaled coefficient [29, 36] to model the remaining unmappable effects:

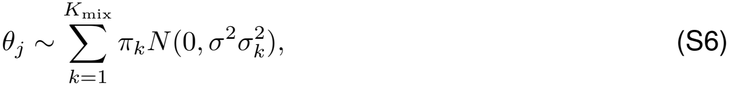

The *K*_mix_ mixture weights 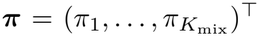 are nonnegative and sum to one. The variance grid is fixed, with 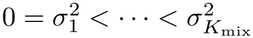.

#### Remark 1 (Comparison with SuSiE and SuSiE-inf)

Standard *SuSiE* models sparse effects (***β***). Both *SuSiE-inf* and *SuSiE-ash* also model background genetic effects (***θ***). *SuSiE-inf* uses a normal prior for these effects, whereas *SuSiE-ash* uses a mixture of normals. In our complex simulations, *SuSiE-inf* had lower power and a lower FDR than standard *SuSiE* (**Supplementary Figure S6**).

### S2 Variational Inference Framework for *SuSiE-ash*

*SuSiE* and *Mr.ASH* use factorized variational approximations [37] and optimize the evidence lower bound (ELBO) by coordinate ascent [38]. In *SuSiE*, iterative Bayesian stepwise selection (IBSS) updates each single effect by fitting SER to partial residuals. *Mr.ASH* updates each background coefficient under a normal mixture prior.

We use a variational distribution that factorizes across the single effects and background coefficients.

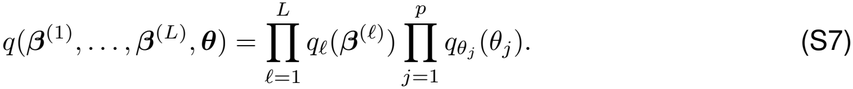

Write 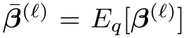 for the posterior mean contribution of single effect *l* and 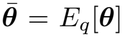 for the posterior mean background effects.

#### Update for the sparse effect *β*

Each single effect is updated by SER using the residual after subtracting the background fit and the posterior mean fitted values of the other sparse effects.

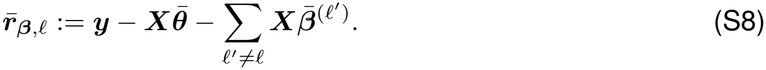

#### Update for the unmappable effects *θ*

We update ***θ*** by fitting *Mr.ASH* to residuals after subtracting the fitted values of sparse effects classified as confident.

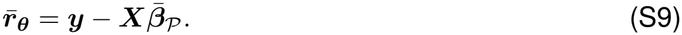

Here 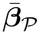 is the sum of the posterior mean coefficients for confident effects (Section S3). We fit *Mr.ASH* to these residuals to estimate 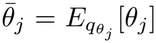 and the residual variance 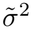. We set 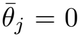 at each masked variant *j* ∈ ℳ.

We alternate the sparse and background updates, checking the single effect inclusion probabilities and PIPs for convergence or a short cycle. Algorithm 1 gives the masking rules. After the loop, the implementation refits the background to residuals that subtract the posterior mean fitted values of all sparse effects.

##### Remark 2 (Bayes factors in Mr.ASH updates)

Let 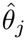 be the working estimate with sampling variance 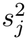, and let 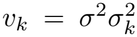 be the prior variance of component *k*. The posterior component weights are *ϕ*_1*jk*_ ∝ *π*_*k*_ BF_*jk*_, where

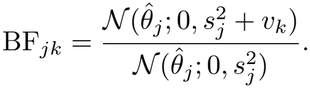

The zero-variance component has BF_*jk*_ = 1 [29].

### S3 Diffusion-Aware Effect Separation

In regions with multiple causal variants in LD, a *SuSiE* single effect can concentrate on a non-causal proxy. We use posterior probabilities and LD to decide where to set the *Mr.ASH* posterior means to zero. For an effect that remains uncertain, we can temporarily lift the mask at its sentinel and nearby variants in tight LD. The next sparse update then uses residuals that account for the background effects estimated at those variants.

#### Effect classification

Let ***α***^(*l*)^ denote the SER inclusion probabilities for effect *l*. For each effect, we form a working credible set 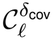 by adding variants in decreasing order of 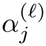 until the cumulative mass exceeds *δ*_cov_ or all variants are included. We write purity_*l*_ for the smallest absolute genotype correlation between distinct members of this set, with purity_*l*_ = 1 for a singleton. The sentinel 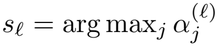 is the variant with the highest inclusion probability for effect *l*. We track effects that have had a sentinel collision or have switched between the uncertain and confident categories in *D*. A collision occurs when two eligible effects have 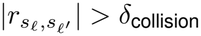. Effects are eligible for this check when the range of their inclusion probabilities is at least 5 × 10^*−*5^.

**Emerging** effects have working credible set purity below *δ*_form_. For these effects we use the sentinel’s moderate LD neighborhood and variants with 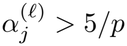 to determine the mask.

**Uncertain** effects have purity at least *δ*_form_ and either have purity below *δ*_pure_ or belong to *D*. We apply `WaitThenExpose` as described below.

**Confident** effects have purity at least *δ*_pure_ and are outside D. We sum their posterior mean coefficients.

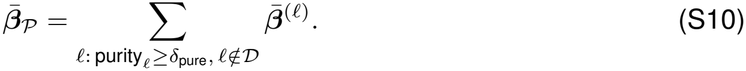

We subtract the fitted values 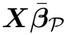 from ***y*** before fitting *Mr.ASH* (Eq. S9).

#### Masking rule

For sentinel *s*_*l*_, define the LD neighborhoods 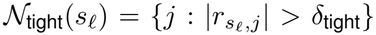 and 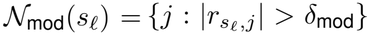. Let *P*_*l*_ be the variants whose probabilities for effect *l* contribute to the mask. For emerging effects, 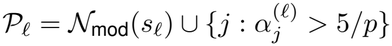. For confident effects, *P*_*l*_ = {1, …, *p*}. For uncertain effects, *P*_*l*_ is empty during a current collision and otherwise includes all variants except any temporarily exposed variants.

We set 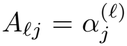 for *j* ∈ *P*_*l*_ and *A*_*lj*_ = 0 elsewhere. The inclusion score used for masking is 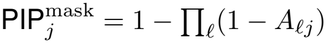. The threshold rule proposes the following variants for masking.

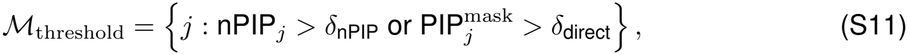

where 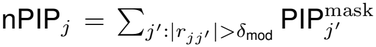. The direct threshold also applies to isolated signals. We add the moderate LD neighborhoods of emerging effects to the proposed mask. The final mask ℳ also accounts for the mask from the previous iteration and temporary exposure.

#### Handling uncertain effects

For an uncertain effect, a current collision resets its waiting counter. A change in sentinel to a variant outside the tight LD neighborhood also resets the counter. After *K* eligible iterations, we temporarily unmask variants in N_tight_(*s*_*l*_) whose masks have yet to be restored after an exposure. After *K*^*′*^ iterations, we restore their masks. For ordinary unmasking, a variant must fall outside the proposed mask for two consecutive iterations.

We keep effects in *D* for the rest of the fit. They remain eligible for credible set reporting. Excluding their fitted values from 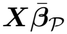 lets the background fit account for residual signal that may arise from several variants in LD.

##### Remark 3 (Default Parameters)

We use 90% posterior mass for working credible sets (*δ*_cov_ = 0.9) and 95% for final reporting by default. The purity cutoffs are *δ*_form_ = 0.1 and *δ*_pure_ = 0.5. We use *δ*_mod_ = 0.5 to define moderate LD and apply direct and neighborhood masking cutoffs of *δ*_direct_ = 0.1 and *δ*_nPIP_ = 0.4. We flag a collision when the absolute genotype correlation between two eligible sentinel variants exceeds *δ*_collision_ = 0.9. Temporary exposure uses the tighter LD threshold *δ*_tight_ = 0.95. We wait *K* = 2 iterations before exposure and *K*^*′*^ = 3 more iterations before restoring the mask.

###### Algorithm 1

Diffusion-Aware Effect Separation for *SuSiE-ash*

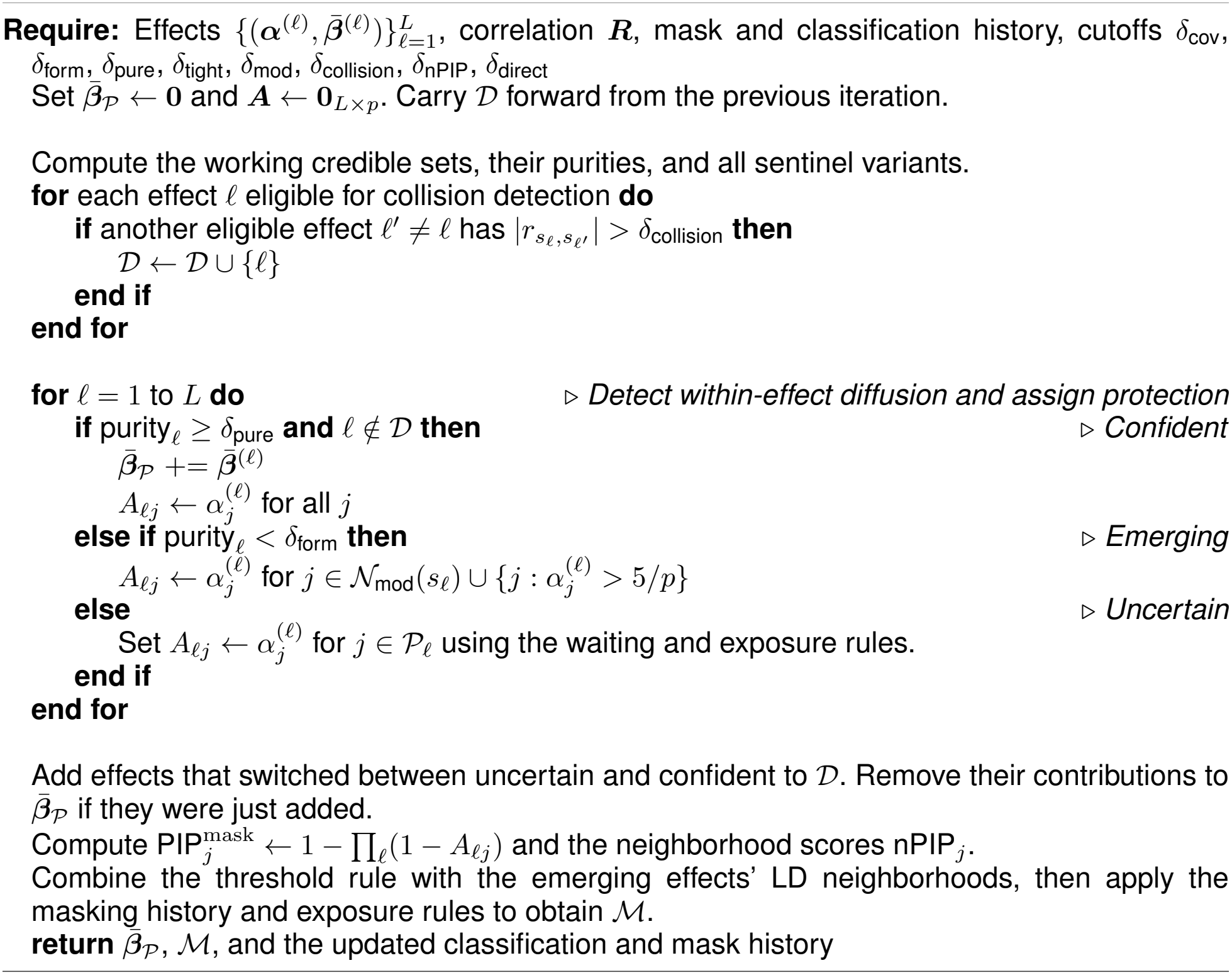

### S4 Implementing *SuSiE-inf* using *Mr.ASH*

*SuSiE-inf* [2] extends *SuSiE* to model infinitesimal effects alongside sparse causal signals by incorporating a random-effects component ***Xθ*** with *θ*_*j*_ ~ *N* (0, *τ* ^2^), leading to the precision matrix

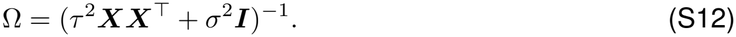

The original implementation estimates *τ* ^2^ via method-of-moments combined with BLUP residuals, which can become computationally intensive for loci with many variants. We also tested an alternative that estimates *τ* ^2^ from the fitted *Mr.ASH* mixture [29].

Similar to *SuSiE-ash*, we model the unmappable background as 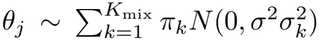 using a fixed variance grid 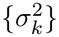 and estimate the mixture weights ***π*** via the *Mr.ASH* coordinate-wise updates. Then, following the *SuSiE-inf* framework, the infinitesimal variance is approximated as prior variance 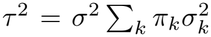, and this estimate is used to construct Ω. For computational efficiency, we use the prior estimate of *τ* ^2^ (from the current mixture weights) when constructing the variance grid at each iteration, rather than re-estimating from posterior moments. We used *SuSiE-inf* to initialize the fit in `susieR` 0.14.55. In version 0.14.56 we used a wider *Mr.ASH* grid and omitted that initialization. We removed this option from later versions because it offered little practical benefit over *SuSiE-inf*.

### S5 Accelerating *SuSiE-inf*

*SuSiE-inf* [2] augments *SuSiE* with a random-effects term, modeling the phenotype as

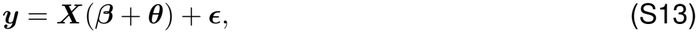

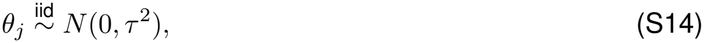

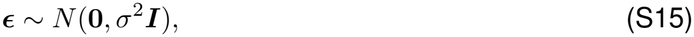

where ***β*** are the sparse single-effect contributions. Absorbing the random effects and noise into an effective residual 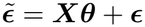 gives the equivalent form

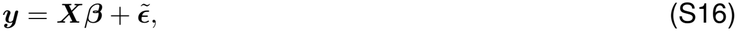

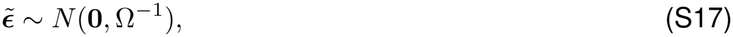

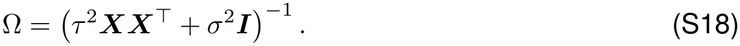

#### Eigendecomposition bottleneck

*SuSiE-inf* carries out its updates using the eigenbasis of the LD matrix ***R*** = ***X***^*⊤*^***X****/n*. A full eigendecomposition of this *p* × *p* matrix has a one-time cost of *O*(*p*^3^). When variants outnumber samples, ***R*** has rank at most *n*, so at most *n* of its *p* eigenvalues are nonzero.

#### Low-rank decompositions

In QTL mapping, variants typically outnumber samples substantially (*p* ≫ *n*). We exploit the resulting low-rank structure in two ways, depending on the available data.

- **Summary-statistics path**. When the LD matrix ***R*** is supplied, we use pivoted Cholesky factorization to obtain an *r* × *p* factor, where *r* is the numerical rank determined by the factorization tolerance. For an LD matrix computed directly from genotypes, the rank is bounded by the number of samples used to estimate LD. The factorization costs *O*(*p*^2^*r*). A thin SVD of this factor gives the eigenvalues and eigenvectors of the rank-*r* approximation to ***R***.
- **Individual-level path**. When the genotype matrix ***X*** is supplied, we take its thin SVD directly. Its right singular vectors are the eigenvectors of ***R***, and its singular values determine the eigenvalues. For *p > n*, this costs *O*(*pn*^2^) and avoids forming or storing the *p* × *p* LD matrix.

The eigenbasis calculations cost *O*(*pr*) per single effect for *r* retained components. For in-sample LD or individual-level genotypes, *r* ≤ *n*. The summary-statistics implementation still uses dense LD matrix-vector products to update residuals and fitted values, so its overall iterative cost remains *O*(*TLp*^2^). With individual-level genotypes, the corresponding iterative cost is *O*(*TLpn*).

#### Complexity

The table summarizes the dominant computational costs when *n* ≪ *p*, where *T* is the number of outer IBSS iterations and *L* is the number of single effects. The decomposition is performed once; the iteration column includes all *T* iterations.

| Method | One-time decomposition | All T IBSS iterations | Total |
|---|---|---|---|
| Original full eigendecomposition | O(p<sup>3</sup>) | O(TLp<sup>2</sup>) | O(p<sup>3</sup> + TLp<sup>2</sup>) |
| LD: pivoted Cholesky + SVD | O(rp<sup>2</sup>) | O(TLp<sup>2</sup>) | O(rp<sup>2</sup> + TLp<sup>2</sup>) |
| Genotypes: thin SVD | O(pn<sup>2</sup>) | O(TLpn) | O(pn<sup>2</sup> + TLpn) |

#### Numerical equivalence

When all discarded eigenvalues are zero, the low-rank decomposition is equivalent to the full eigendecomposition in exact arithmetic. Across simulated replicates, the accelerated methods matched the original posterior inclusion probabilities and variance components to machine precision and returned identical credible sets.

### S6 Accelerating the Regularized RSS Model

The regularized RSS model [25] fits *SuSiE* to summary statistics using the residual covariance **Σ** = *σ*^2^***R*** + *λ****I***, where ***R*** is the LD matrix and *λ* is a positive ridge parameter that keeps **Σ** invertible when ***R*** is rank deficient. Here, the acceleration comes from avoiding repeated dense matrix calculations when evaluating the objective and updating the residual variance.

Both implementations decompose ***R*** once. The original implementation repeatedly formed dense matrix products to evaluate quadratic forms and traces during objective evaluation and residual-variance estimation. *SuSiE 2.0* evaluates these quantities using matrix-vector products in the eigenbasis and sums over eigenvalues. A dense transform is still used for the single-effect updates. These changes preserve the model and update equations while reducing the cost of these calculations.

Benchmarks used *L* = 5, *λ* = 10^*−*5^, and 100 replicates per setting, with speedups calculated as ratios of median runtimes. Relative to the original `susie_rss_lambda` implementation, the summary-statistics path achieved a 7.0-8.5-fold speedup on identical inputs across variant-to-sample ratios from 5 to 50 at *p* = 5,000. Supplying a genotype matrix instead of a dense LD matrix increased the speedup to 11.2-16.6-fold, with larger gains as variants outnumbered samples (**Supplementary Figure S1**).

### S7 Correlation-Based Credible Set Extension

A 95% credible set is a set of variants that jointly carry at least 95% of the posterior mass of a single effect. When several variants are in near-perfect LD, posterior mass can be divided among them, leaving highly correlated proxies outside the reported set. *SuSiE 2.0* therefore provides an optional correlation-based extension, controlled by the `cs_extension_corr` argument of `susie_get_cs`. Given a threshold *ρ* ∈ [0, 1], each credible set is extended to include every variant whose absolute correlation with at least one member of the original set exceeds *ρ*. The posterior mass is then recomputed over the extended set, and the extended set remains subject to the usual purity filter. The extension can use either a dense genotype matrix ***X*** or a correlation matrix. It is disabled by default; when used, the recommended threshold is *ρ* = 0.99, restricting additions to variants in very high LD with an existing member.

At *ρ* = 0.99, we observed small improvements in power and FDR for *SuSiE, SuSiE-ash*, and *SuSiE-inf* under sparse and complex architectures (**Supplementary Figures S8 and S9**). Extension can recover a causal variant omitted in favor of a highly correlated proxy, although improvements are not guaranteed for every data set.

### S8 Simulation Study Details

#### Sparse effects simulation design

We conducted extensive simulations to evaluate the performance of *SuSiE, SuSiE-ash*, and *SuSiE-inf* under a sparse genetic architecture, with slight modifications to the original *SuSiE* paper’s simulation design. These simulations used genotype data from UK Biobank where we randomly sampled 150 LD blocks across chromosomes 1-22 and derived the corresponding genotype matrices to serve as the basis for our simulation. Each genotype matrix, **X** ∈ ℝ^*n×p*^, contained *n* = 1,000 individuals and at most *p* = 5,000 variants after filtering. Additionally, we required a minor allele frequency greater than 1%, and a missing rate below 5% (using mean imputation for missing data).

We generated gene expression levels, **y** ∈ ℝ^*n*^, based upon the following linear model:

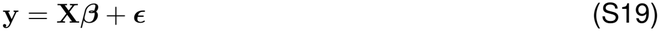

where ***β*** ∈ ℝ^*p*^ denotes the effect size vector and ***ϵ*** ∈ ℝ^*n*^ denotes the noise vector, such that ***ϵ*** ~ *N* (0, *σ*^2^**I**_*n*_) where **I**_*n*_ is the *n* × *n* identity matrix. The effect size vector, ***β*** ∈ ℝ^*p*^, was constructed by sampling a set of causal variant indices, S, uniformly at random, subject to pairwise absolute genotype correlations of at most 0.1. For each causal variant, *j* ∈ *S*, we set *β*_*j*_ = 1, and for all non-causal variants, *j* ∉ *S*, we set *β*_*j*_ = 0. Finally, to achieve the desired heritability-level, *h*^2^, we set the residual variance, *σ*^2^, to solve the following equation:

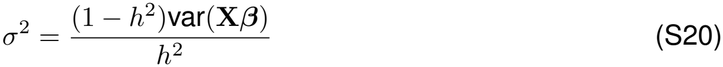

For each simulated effect, we fixed the per-SNP heritability 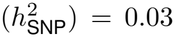. We then generated simulated data sets varying the number of causal variants *k* = {1, 2, 3, 4, 5} resulting in *h*^2^ = {0.03, 0.06, 0.09, 0.12, 0.15}, respectively. For each of the five scenarios, we generated 500 replicates for fine-mapping and 300 replicates for prediction.

#### Complex genetic architecture simulation design

We also evaluated *SuSiE, SuSiE-ash*, and *SuSiE-inf* under multiple complex genetic architectures to mimic a realistic eQTL architecture [26]. In these settings, gene expression is influenced both by a core set of variants that exert medium to large effects and by a polygenic background in which i) a number of variants (15) each contribute a small amount of phenotypic variance, or ii) all remaining variants each contribute a non-zero amount of variation. We used the same preprocessed genotype matrices from the UK Biobank data described in the sparse simulation design section.

We generated gene expression levels, **y** ∈ ℝ^*n*^, using a linear model that partitions genetic effects into three distinct components: sparse, oligogenic, and polygenic. The sparse component, *S*, includes *k* = 3 variants which exhibit relatively large effects. These effects are drawn from a normal distribution, 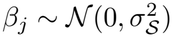, and then scaled to the nominal genetic variance target 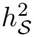.

The oligogenic component, *O*, has a small number of variants (5-10) that contribute moderate effects. These effects are modeled using a two-component Gaussian mixture distribution to allow for added variability in their magnitudes:

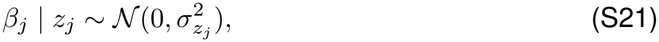

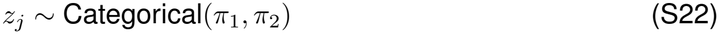

for *j* ∈ *O*, where *z*_*j*_ ∈ {1, 2} denotes the mixture component assignment, 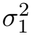 and 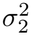 are the component-specific variances with 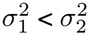 and *π*_1_ +*π*_2_ = 1. The oligogenic effects are then scaled to the nominal genetic variance target 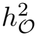.

The polygenic component, *P*, models the contribution of remaining causal variants. Effects for this component are drawn from a normal distribution with small variance, 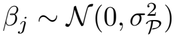 for *j* ∈ *P*, and scaled to the nominal genetic variance target 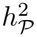. Variants not assigned to any component have effects set to zero.

For complex architectures, we initially required pairwise absolute genotype correlations below 0.1 among the sparse and oligogenic variants, including pairs from different components. After 500 unsuccessful sampling attempts at a given threshold, we increased the threshold by 0.1, up to 0.5. If no configuration satisfied these restrictions, we used the last one sampled. No corresponding LD restriction was imposed on the polygenic variants.

Collectively, these three mutually exclusive sets partition the nonzero-effect variants, so that *S* ∪ *O* ∪ *P* = {*j* : *β*_*j*_≠ 0}. These components comprise the effect vector ***β*** ∈ ℝ^*p*^ to create the following linear model:

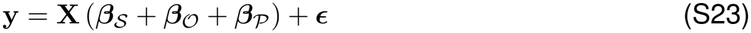

where **X** ∈ ℝ^*n×p*^ is the standardized genotype matrix and ***ϵ*** ∈ ℝ^*n*^ is a noise vector with ***ϵ*** ~ *N* (0, *σ*^2^**I**_*n*_) where **I**_*n*_ is the *n* × *n* identity matrix.

Each component was scaled separately to its nominal genetic variance target. We then set the residual variance from the variance of the combined genetic contribution to achieve the desired total heritability. Because the components can be correlated, their realized heritability contributions may differ from the nominal proportions specified below.

In our benchmark, we evaluated three scenarios with varying genetic architectures, all with total heritability 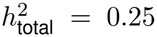, using 500 replicates per scenario for fine-mapping and 300 for prediction. The first scenario, oligogenic effects on a polygenic background, included 3 sparse effects (50% of 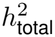), 5 oligogenic effects (35% of 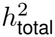), and 15 polygenic effects (15% of 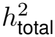). The second scenario, oligogenic effects on a *moderate* infinitesimal background, maintained the same heritability proportions (50%, 35%, 15%) and variant counts for sparse and oligogenic components, but assigned the remaining heritability to *all* remaining variants. The third scenario, oligogenic effects on an *extensive* infinitesimal background, shifted the architecture toward a stronger infinitesimal contribution: 3 sparse effects (50% of 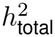), 10 oligogenic effects (15% of 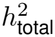), and all remaining variants (35% of 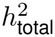).

For complex architectures, we evaluated fine-mapping performance using the *N* variants with the largest absolute nonzero simulated effects as the target set, with *N* ∈ {3, 5, 10, 15, 20, 25}. When fewer than *N* variants had nonzero effects, all nonzero-effect variants were included. Power was the proportion of target variants contained in at least one reported CS. The false discovery proportion was the proportion of reported CSs containing none of the target variants. We averaged these quantities across replicates to obtain power and FDR, assigning both quantities a value of zero when no CSs were reported.

For reference, the figures reporting each complex scenario’s results are summarized below.

| Scenario | Fine-mapping (power/FDR) | TWAS ($R^2$) | CS size | Illustrative locus | Runtime |
|---|---|---|---|---|---|
| Polygenic background | Figure S9E, S9F | Figure S4B, S4F | Figure S3B | Figure S5 | — |
| Moderate infinitesimal | Figure 1C, 1D, S6A, S6B, S9C, S9D | Figure S4C, S4G | Figure S3C | — | — |
| Extensive infinitesimal | Figure 1E, 1F, S6C, S6D, S9A, S9B | Figure S4D, S4H | Figure S3D | — | Figure S11 |

### S9 Evaluation of Alternative Credible Set Coverage Methods

We evaluated attainable coverage and Bayesian Linear Programming (BLiP) [27] as alternatives to standard SuSiE credible set reporting (**Supplementary Figure S7**).

When LD is unavailable for purity filtering, one can use the attainable coverage criterion introduced by SparsePro to screen effects. The original SparsePro publication [21] describes constructing sets at the requested coverage for effects that pass this screen. At the nominal 95% level, the released code targets 0.95*A*, where *A* is the attainable mass rounded to two decimal places. It can also stop sooner when the next variant has weight below 0.01. A reported set can therefore contain less than 95% of the original single-effect posterior. In *SuSiE 2.0*, we construct each reported set from the full single-effect posterior at the requested coverage. For the 567 effects reported by both rules in simulations using a 121-variant ADSP LD window, 152 Sparse-Pro sets contained less than 95% of the original posterior mass. All corresponding *SuSiE 2.0* sets reached 95% (**Supplementary Figure S7E**).

The SparsePro default entropy threshold of log(20) performed poorly in both our sparse and complex simulations, which used dense marker sets (*p* = 5,000). We use log{max(100, 0.10*p*)} in *SuSiE 2.0*, where *p* is the number of variants. BLiP offered limited gains over default *SuSiE* in the scenarios we evaluated, so we did not include it in the current release.

### S10 Functional Enrichment of Credible Sets

For each method, we extracted all variants within 95% CSs along with their PIPs. Each unique variant was scored using AlphaGenome. We selected tissue-matched AlphaGenome prediction tracks for each brain region: RNA-seq and DNase from dorsolateral prefrontal cortex (UBERON:0009834) for DLPFC, RNA-seq from anterior cingulate cortex (UBERON:0009835) for AC, and DNase from posterior cingulate gyrus (UBERON:0002740) for PCC.

For each variant and modality, we averaged the signed quantile-normalized AlphaGenome scores across tissue-matched tracks before taking the absolute value. RNA-seq scores were additionally matched to the target gene. We then defined the PIP-weighted score of a credible set *C* as

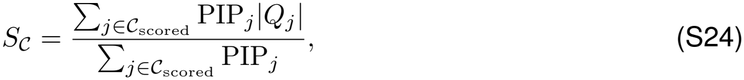

where *Q*_*j*_ is the mean signed score for variant *j*, and *C*_scored_ contains the CS members with available scores for that modality. Credible sets without any scored variants were omitted for that modality.

We restricted the comparison to gene-tissue pairs with results from all three methods where standard *SuSiE* identified at least two CSs. Within each gene-tissue pair, we linked CSs from different methods when their variant-membership Jaccard index was at least 0.75. CSs connected directly or through other linked CSs were grouped as one signal. Matching was performed separately for the two-method comparison of *SuSiE* and *SuSiE-inf* and the three-method comparison that also included *SuSiE-ash*. The two-method comparison produced three groups: shared signals and signals unique to either method. The three-method comparison produced seven groups: signals shared by all three methods, three groups shared by exactly two methods, and three method-unique groups. For each signal and modality, we averaged the available scores of its constituent CSs, so that each signal contributed one observation per modality.

For each group and modality, we used a one-sided *t*-test to assess whether the mean signal score exceeded a reference value of 0.5. We applied the Benjamini-Hochberg correction separately across the six tests in the two-method comparison and the fourteen tests in the three-method comparison.

For the *LMOD1* locus, we additionally summarized RNA-seq support using the maximum absolute score across CS members, max |*Q*_RNA_|. We assessed overlap with ENCODE regulatory annotations obtained through FILER using the original hg38 genomic intervals.

## Supplementary Figures

**Fig. S1:**
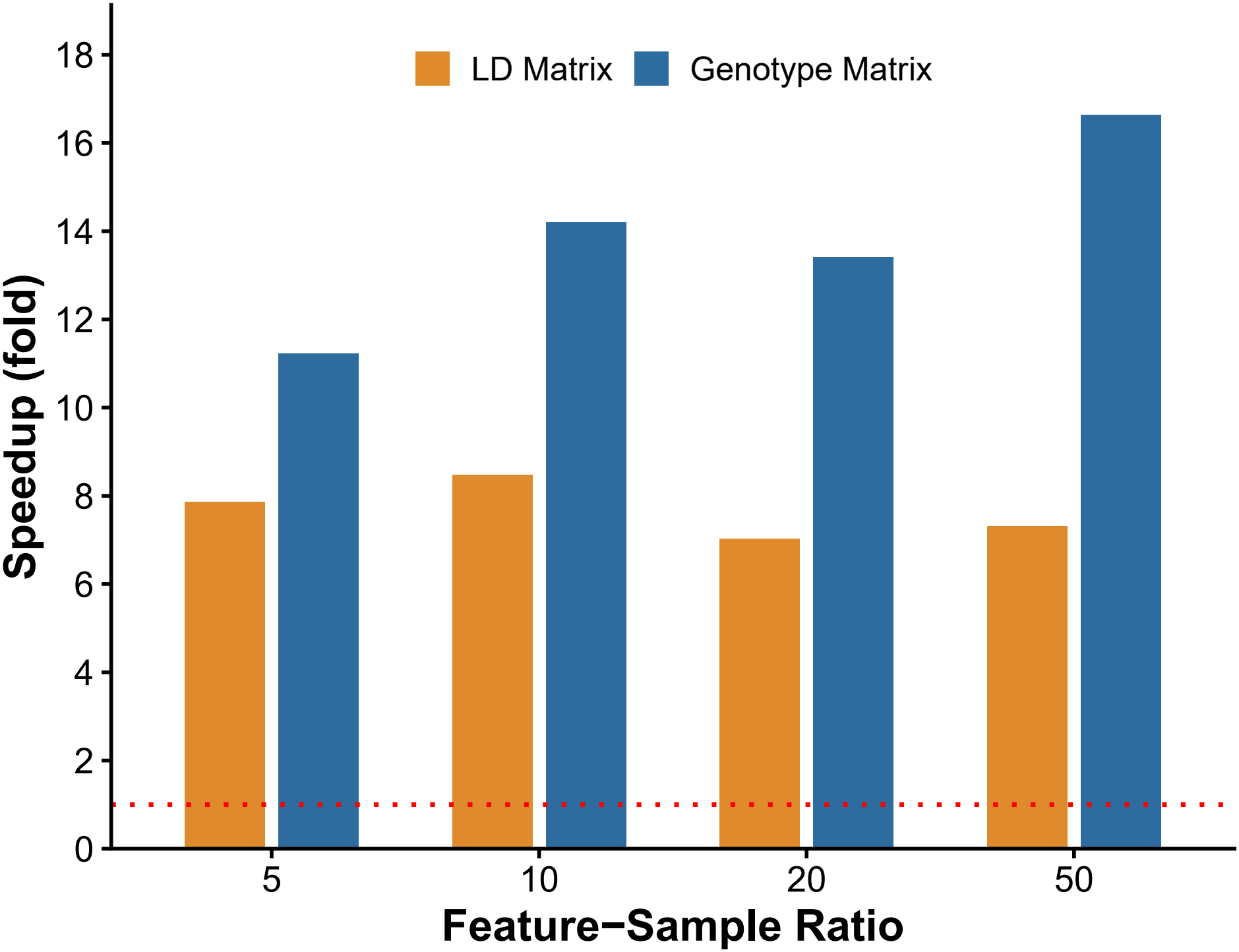
Summary statistics backend speedup for the *λ*-regularized RSS model. Fold speedup of the *SuSiE 2.0* summary statistics backend over the original susieR susie_rss_lambda implementation, using an LD correlation matrix (orange) or a genotype matrix (blue), at *p* = 5,000 across sample sizes.

**Fig. S2:**
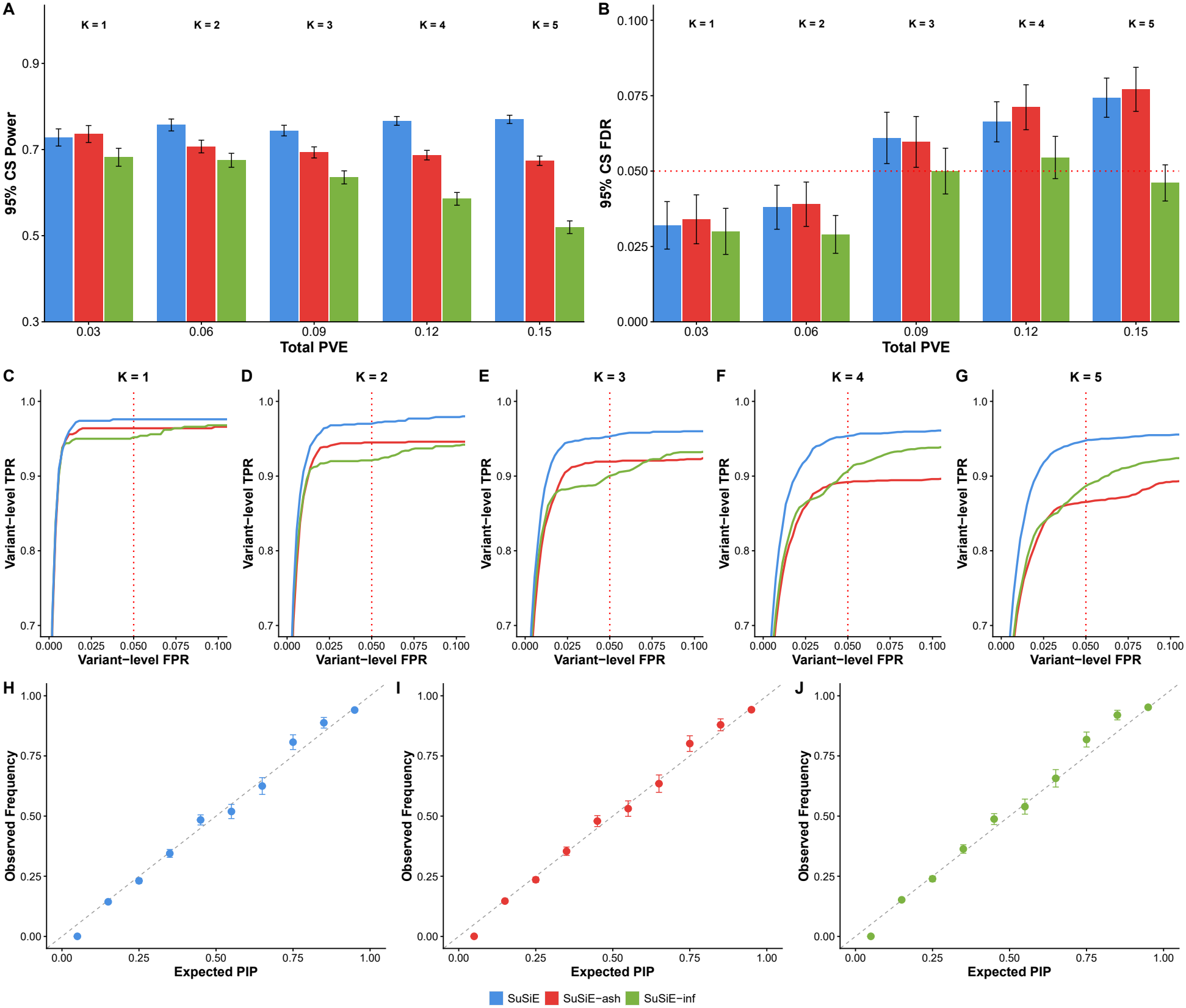
Fine-mapping performance under sparse settings. Comparison of *SuSiE* (blue), *SuSiE-ash* (red), and *SuSiE-inf* (green) using UK Biobank genotypes, with *n* = 1,000, at most 5,000 variants, and 500 replicates for each *K* = 1, …, 5, at fixed per-SNP heritability 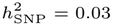. (A, B) Mea 95% CS power and FDR versus total PVE, with error bars showing *±* 1 standard error across replicates. (C-G) Variant-level TPR versus FPR, pooling variants across replicates separately for each *K*. (H-J) PIP calibration for *SuSiE, SuSiE-ash*, and *SuSiE-inf*, respectively, pooling all five settings. Points show observed causal frequencies within PIP bins of width 0.1, plotted at bin midpoints.

**Fig. S3:**
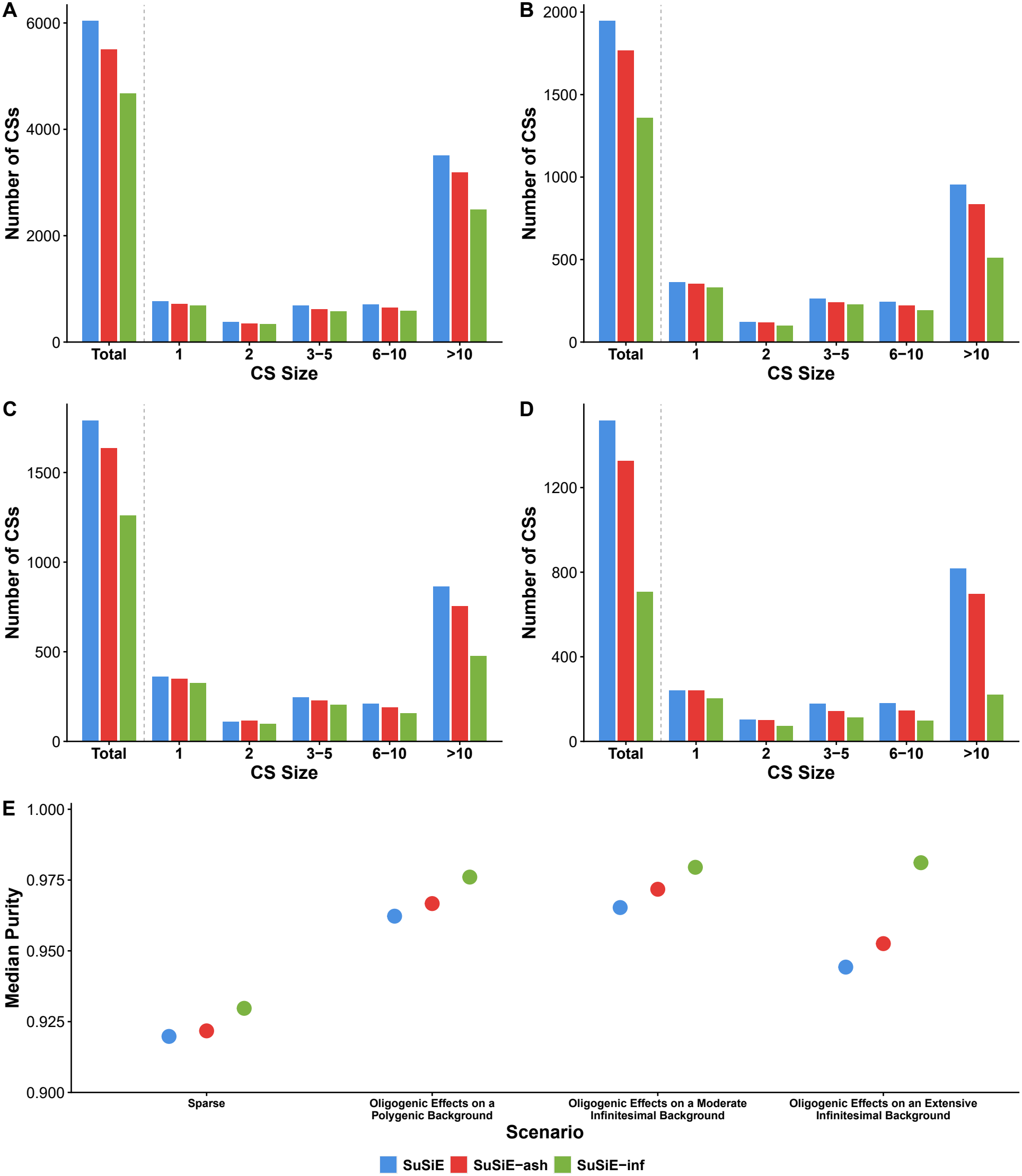
Credible set size distributions and median purity across simulation scenarios. Per-method counts of credible sets binned by size for (A) sparse, (B) oligogenic effects on a polygenic background, (C) oligogenic effects on a moderate infinitesimal background, and (D) oligogenic effects on an extensive infinitesimal background, with (E) median purity across all four scenarios.

**Fig. S4:**
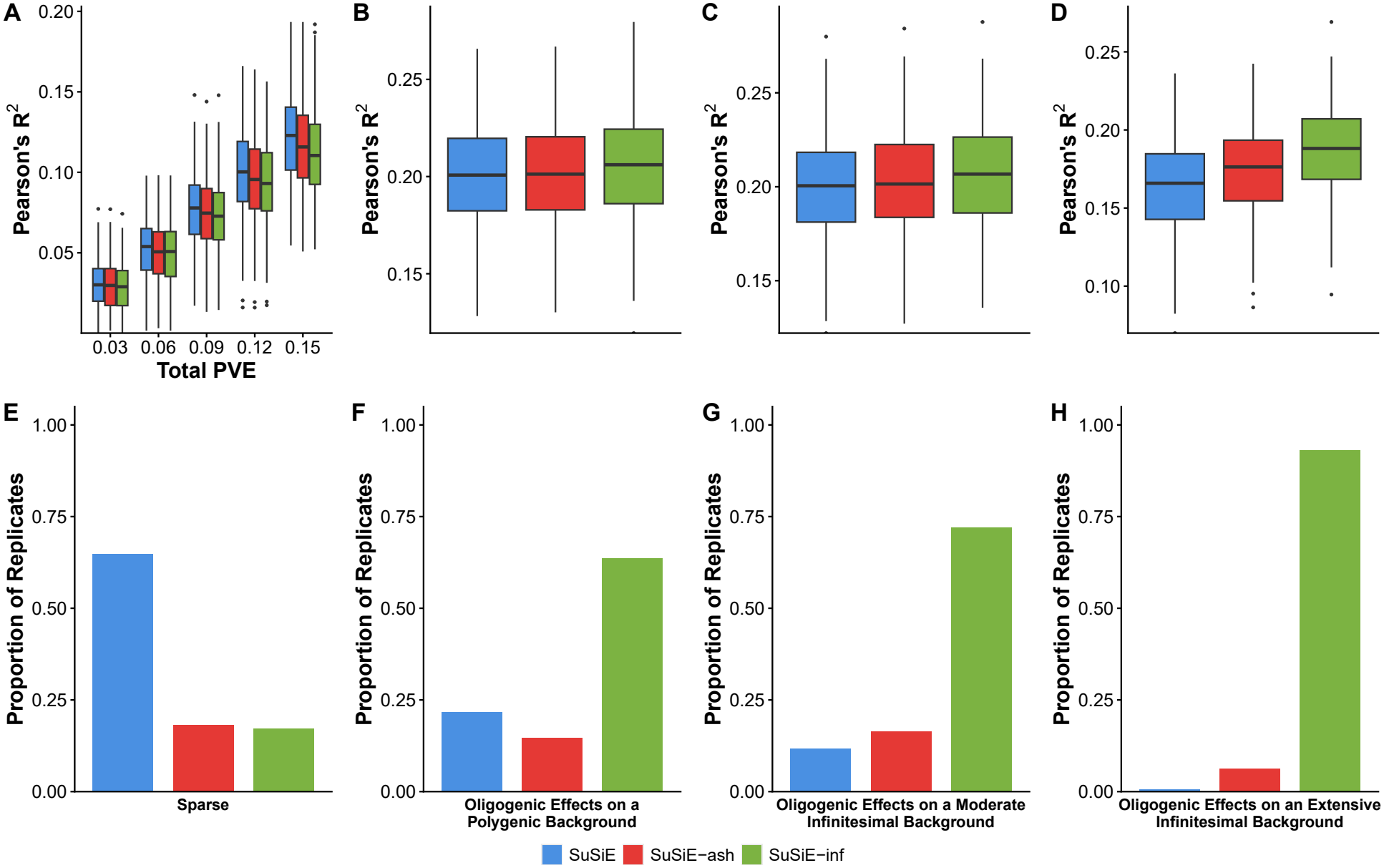
Gene expression prediction in simulations. Cross-validated prediction accuracy (Pearson’s *R*^2^) for *SuSiE* (blue), *SuSiE-ash* (red), and *SuSiE-inf* (green) across simulated genetic architectures, using 300 replicates per setting. (A-D) Distribution of *R*^2^ across replicates. (E-H) Proportion of replicates in which each method achieved the highest *R*^2^.

**Fig. S5:**
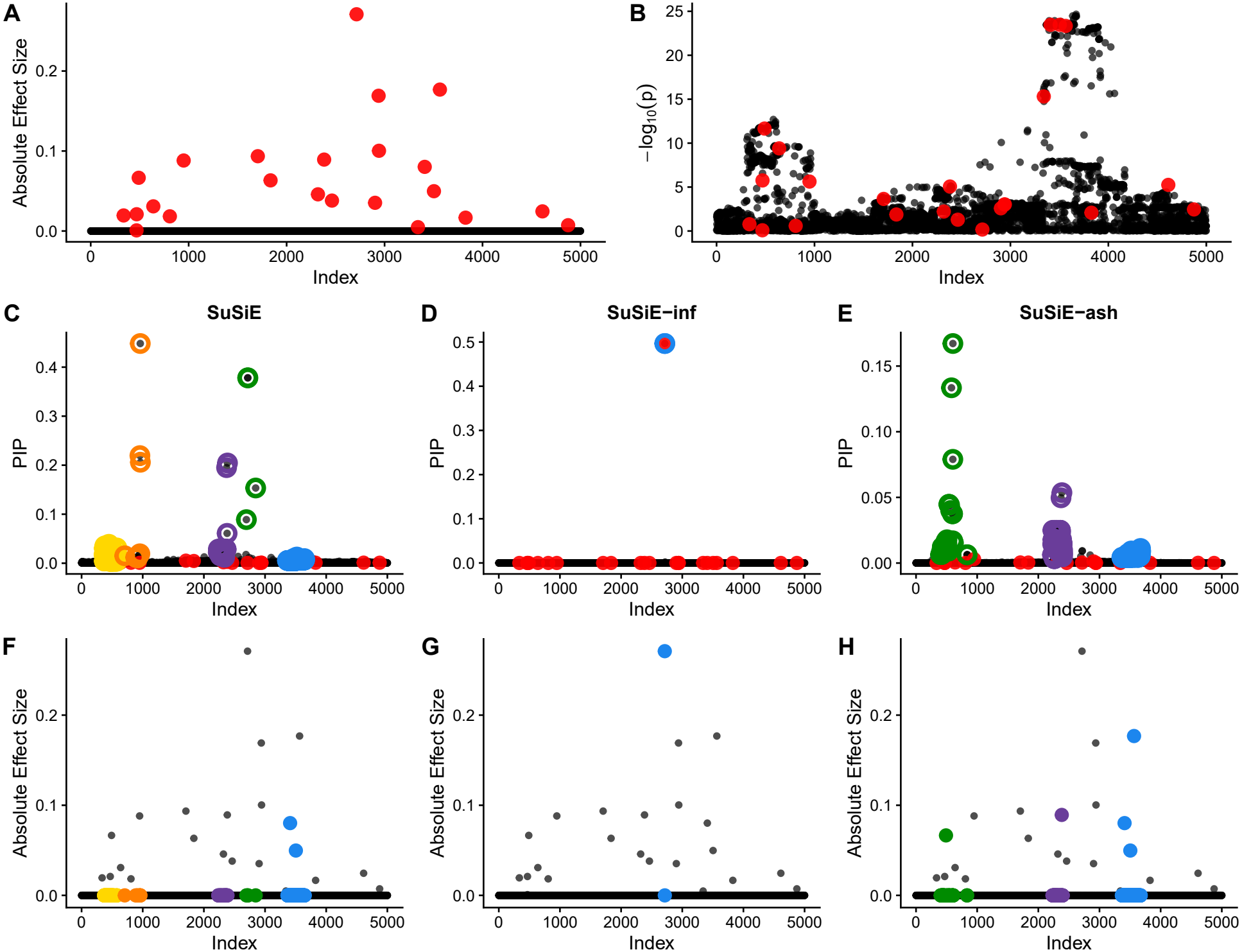
Illustrative locus under polygenic architecture. Comparing *SuSiE, SuSiE-inf* and *SuSiE-ash* on a single simulated locus (*p* = 5,000), showing (A) true absolute effect sizes and (B) marginal association (*−* log_10_ *p*), with causal variants in red, alongside per-variant PIPs (C-E) and credible set assignments (F-H).

**Fig. S6:**
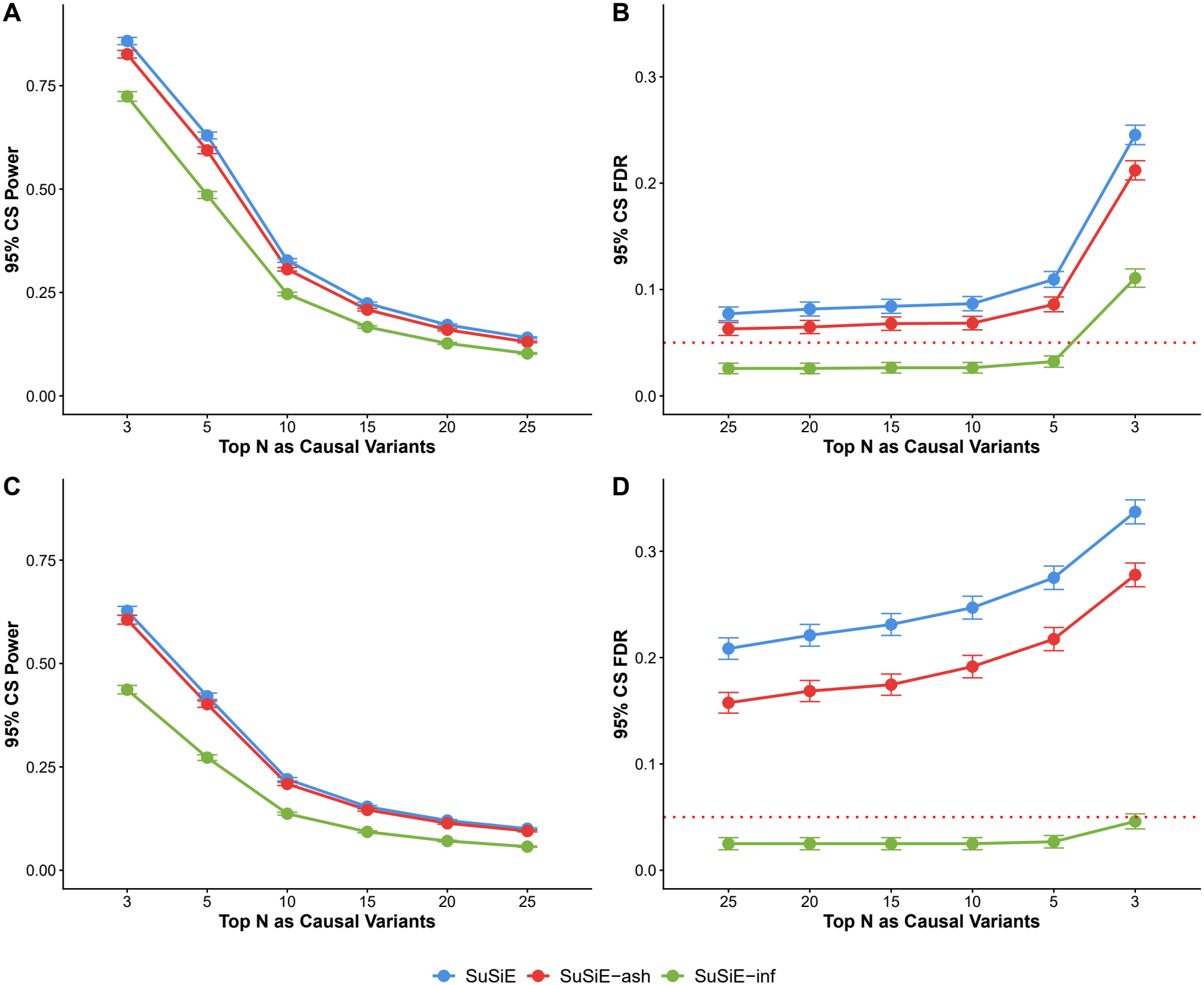
Fine-mapping performance under complex settings, including *SuSiE-ash*. Three-method version of **Figure 1C-F**. (A, C) 95% CS power and (B, D) 95% CS FDR for *SuSiE, SuSiE-ash*, and *SuSiE-inf*, as the number of top variants treated as causal varies, under oligogenic effects on (A, B) a moderate infinitesimal background and (C, D) an extensive infinitesimal background.

**Fig. S7:**
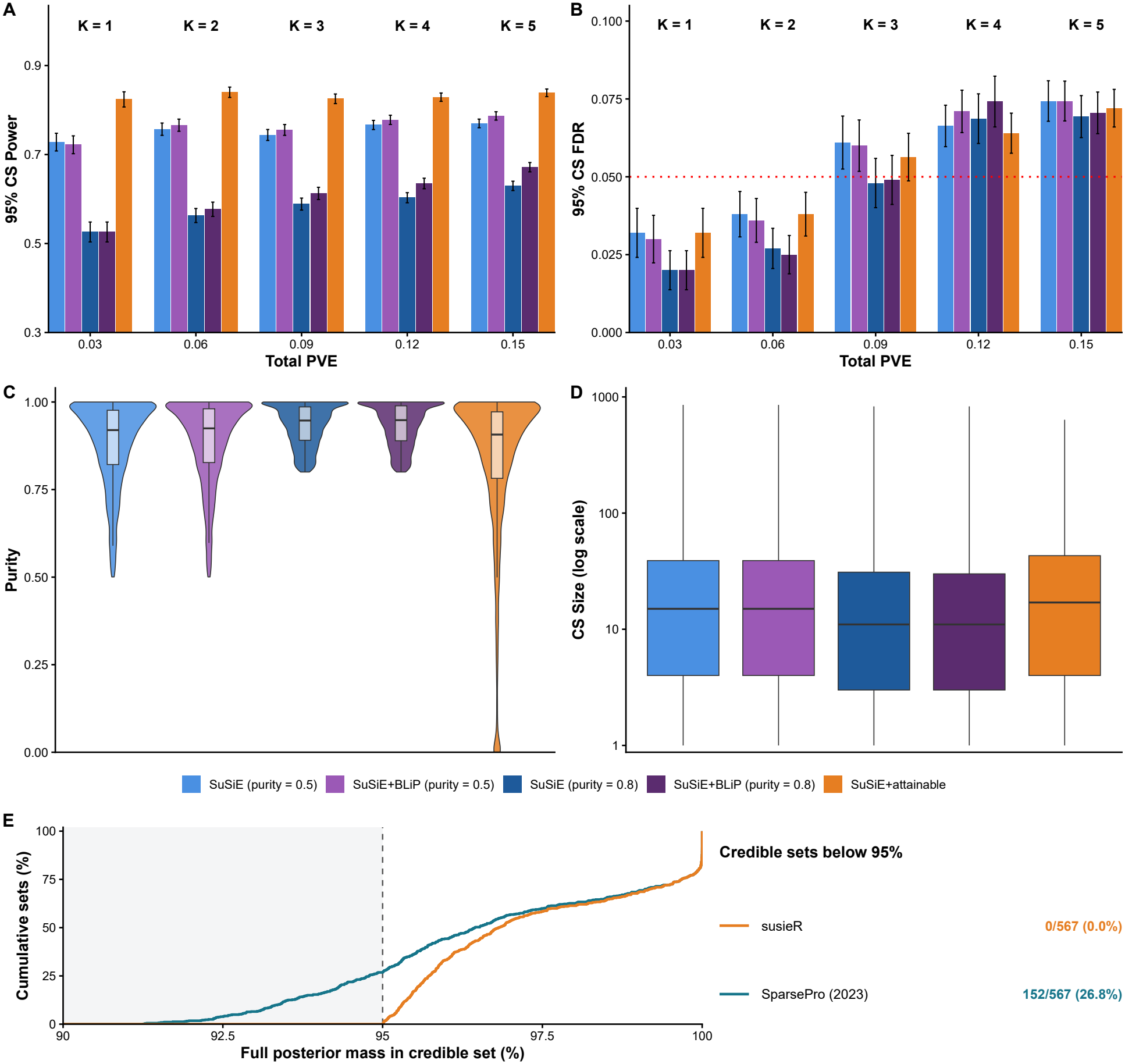
Evaluation of credible set construction strategies under sparse settings. Comparison of *SuSiE* and *SuSiE* +BLiP at purity thresholds of 0.5 (default) and 0.8, and *SuSiE* with attainable coverage. (A) 95% CS power, (B) 95% CS FDR, (C) CS purity, and (D) CS size on a log scale. (E) Cumulative distributions of posterior mass in the credible sets reported by SparsePro and susieR in simulations using ADSP LD. We included fitted effects for which both rules reported a set and measured each set against the full single-effect posterior. The dashed line marks 95%.

**Fig. S8:**
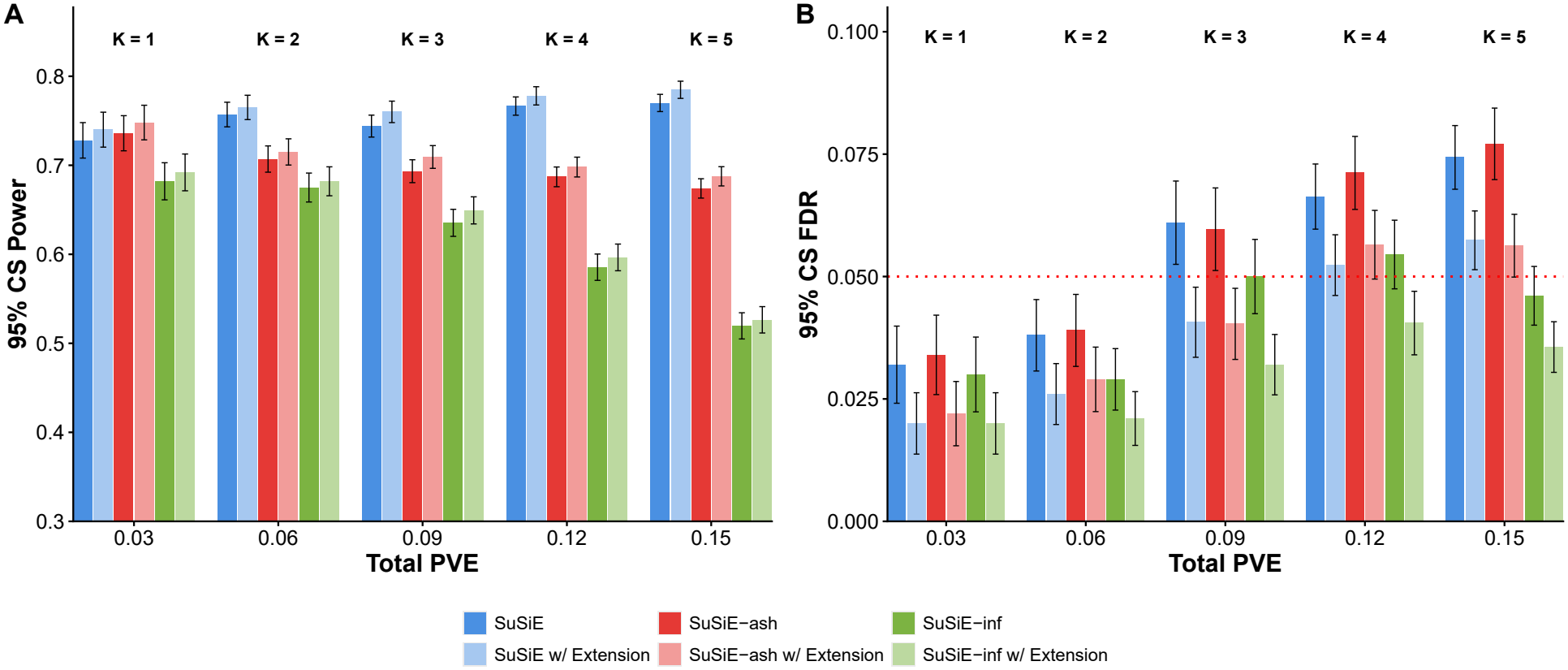
Correlation-based credible set extension under sparse settings. (A) 95% CS power and (B) 95% CS FDR for *SuSiE, SuSiE-ash*, and *SuSiE-inf*, each with and without the extension, across total PVE and number of causal variants (*K* = 1 to 5).

**Fig. S9:**
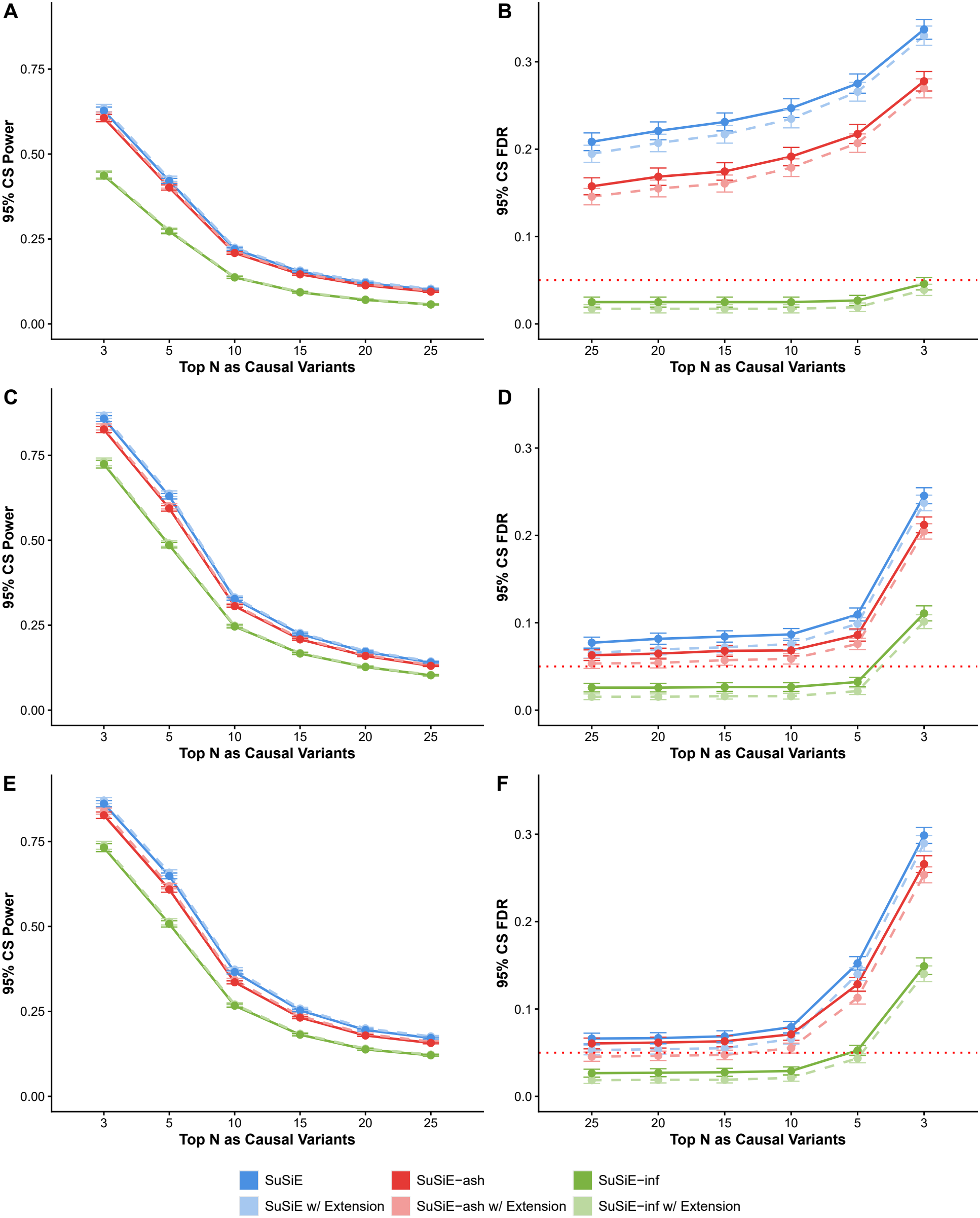
Correlation-based credible set extension under complex settings. (A, C, E) 95% CS power and (B, D, F) 95% CS FDR for *SuSiE, SuSiEash*, and *SuSiE-inf*, each with and without the extension, as the number of top variants treated as causal varies, under oligogenic effects on (A, B) an extensive infinitesimal background, (C, D) a moderate infinitesimal background, and (E, F) a polygenic background.

**Fig. S10:**
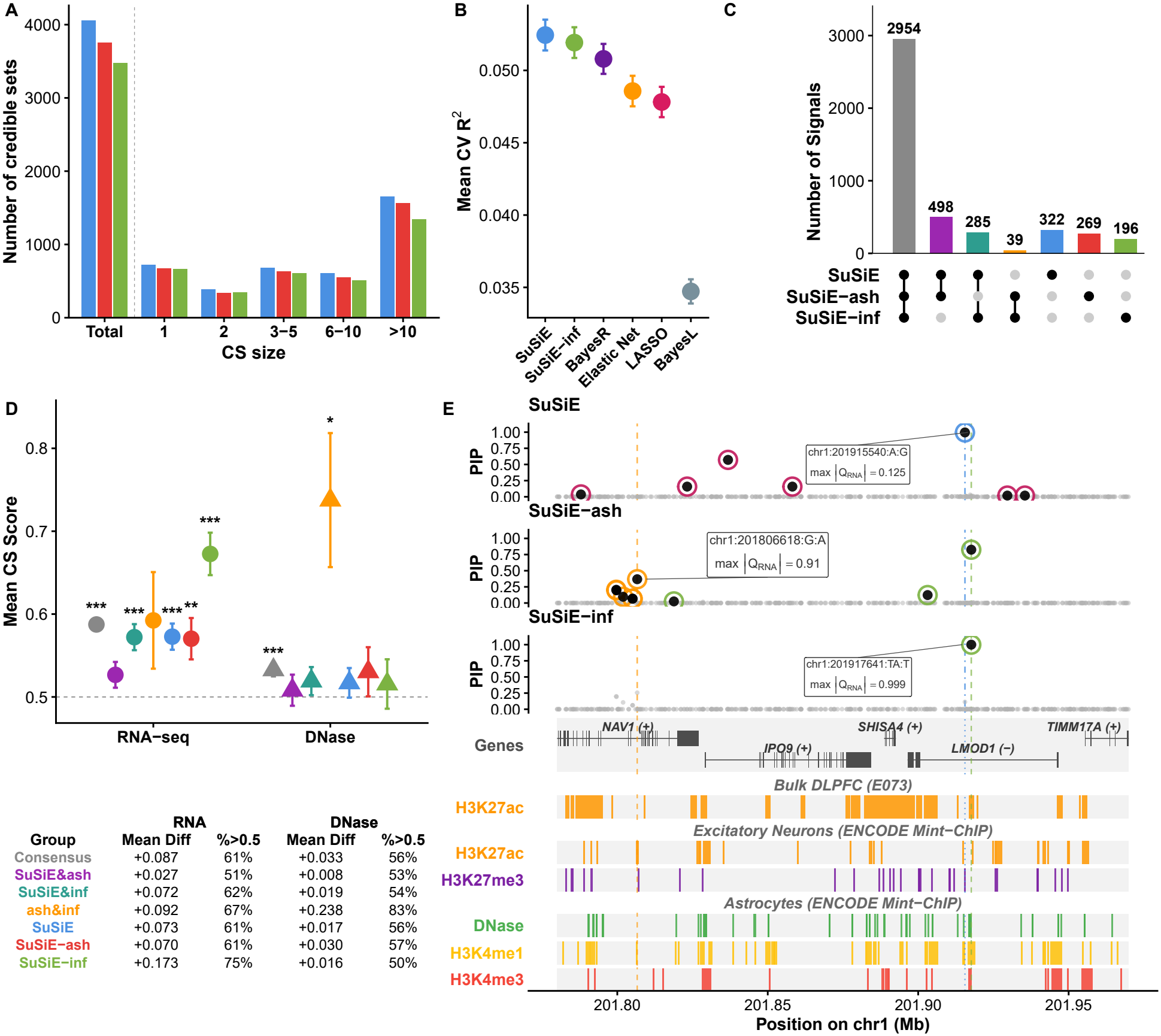
ROSMAP analysis including *SuSiE-ash*. Three-method version of **Figure 2**. (A) Number of credible sets identified by each method, in total and stratified by credible set size. (B) Mean cross-validated *R*^2^ for gene expression prediction methods. (C) Number of signals recovered by all three methods, by a given pair of methods, or unique to one method. (D) PIP-weighted AlphaGenome functional enrichment scores by cross-method concordance group, for RNA-seq (circles) and DNase (triangles), restricted to complex regions where *SuSiE* reports two or more CSs, with points, error bars, and asterisks as in **Figure 2D**; the table gives the mean difference from the 0.5 reference value and the percentage of signals exceeding it. (E) Fine-mapping at the *LMOD1* locus for all three methods, showing per-method PIPs and tissue-matched regulatory annotation tracks.

**Fig. S11:**
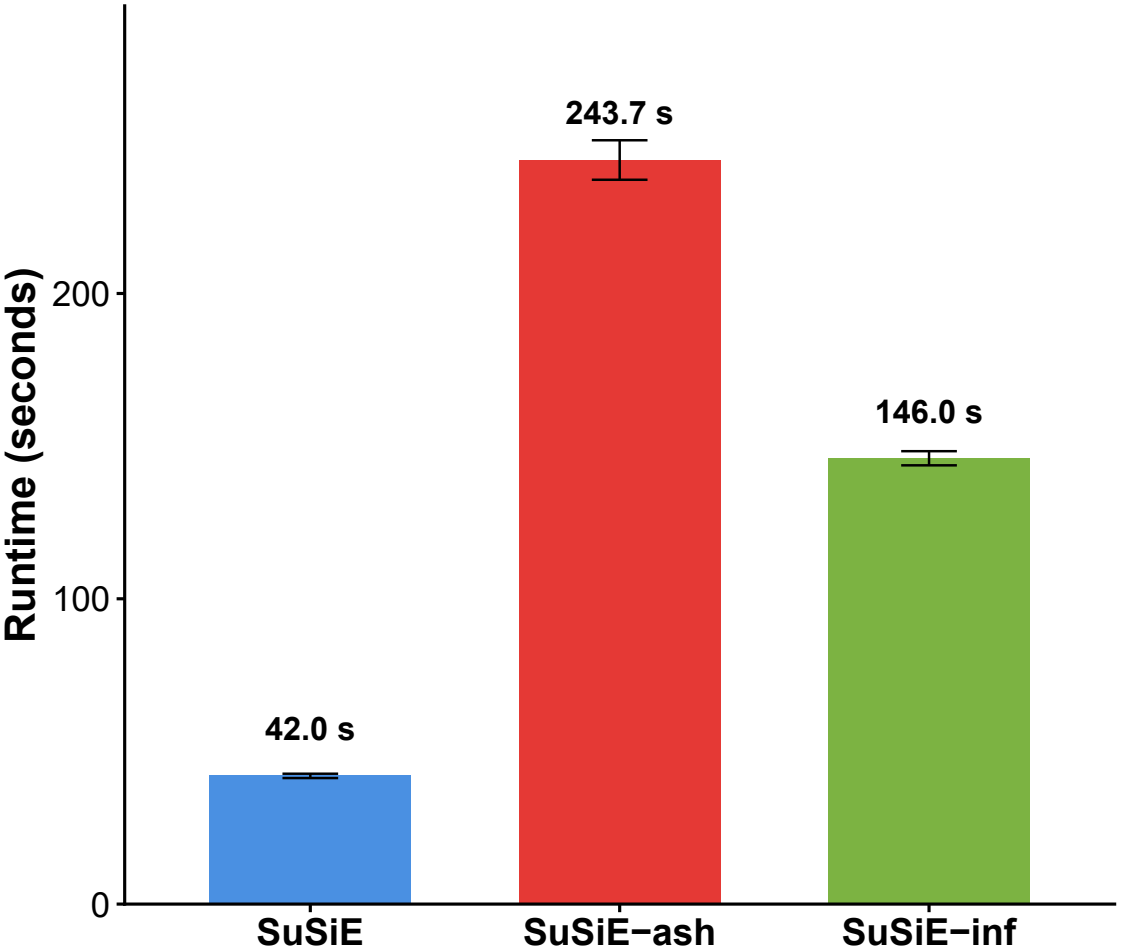
Runtime comparison across methods. Mean runtime comparing *SuSiE* methods over 500 replicates under oligogenic effects on an extensive infinitesimal background (*n* = 1,000, *p* = 5,000), with error bars showing *±* standard error of the mean.

